# Genome-wide characterization of differential transcript usage in *Arabidopsis thaliana*

**DOI:** 10.1101/136770

**Authors:** Dries Vaneechoutte, April R. Estrada, Ying-Chen Lin, Ann E. Loraine, Klaas Vandepoele

## Abstract

Alternative splicing and the usage of alternate transcription start- or stop sites allows a single gene to produce multiple transcript isoforms. Most plant genes express certain isoforms at a significantly higher level than others, but under specific conditions this expression dominance can change, resulting in a different set of dominant isoforms. These events of Differential Transcript Usage (DTU) have been observed for thousands of *Arabidopsis thaliana*, *Zea mays* and *Vitis vinifera* genes and have been linked to development and stress response. However, the characteristics of these genes, nor the implications of DTU on their protein coding sequences or functions, are currently well understood. Here we present a dataset of isoform dominance and DTU for all genes in the AtRTD2 reference transcriptome based on a protocol that was benchmarked on simulated data and validated through comparison with a published RT-PCR panel. We report DTU events for 8,148 genes across 206 public RNA-Seq samples and find that protein sequences are affected in 22% of the cases. The observed DTU events show high consistency across replicates and reveal reproducible patterns in response to treatment and development. We also demonstrate that genes with different evolutionary ages, expression breadths, and functions show large differences in the frequency at which they undergo DTU and in the effect that these events have on their protein sequences. Finally, we showcase how the generated dataset can be used to explore DTU events for genes of interest or to find genes with specific DTU in samples of interest.

**SIGNIFICANCE STATEMENT:** Differential transcript usage through alternative splicing has been reported for thousands of genes in plants, yet genome-wide datasets to study the implications for gene functions are thus far not available. Here we present the first reference dataset of isoform dominance and differential transcript usage for *Arabidopsis thaliana* based on 206 public RNA-Seq samples and provide insights in the occurrence and functional consequences of alternative splicing.

## INTRODUCTION

When and where a gene is transcribed in a eukaryotic cell is governed by a complex system of regulatory elements. Trans-acting transcription factors regulate transcription by binding to cis-regulatory elements on the DNA and by recruiting the transcriptional machinery, including the RNA polymerase enzyme (Hobert, 2008). Access to these cis-regulatory elements depends on the chromatin structure, which is in turn regulated by histone modifications and DNA methylation (Li, 2002). Usage of alternative transcription start- and stop sites and co-transcriptional alternative splicing (AS) of the precursor RNA allow for an assortment of different transcripts or isoforms to be created from a single gene. AS is regulated by transacting splice factors, such as serine/arginine-rich proteins and heterogeneous nuclear ribonucleoproteins, that bind to splice sequences at the intron-exon border or to splicing enhancer- or silencer sequences within introns or exons (Pertea et al., 2007; Erkelenz et al., 2013). Four major types of alternative splice events exist that can change the mRNA content: (i) exon skipping, which is the most frequent in vertebrates and invertebrates (Kim et al., 2007), (ii) intron retention, which is the most frequent in plants (Wang and Brendel, 2006; Filichkin et al., 2010; Marquez et al., 2012; Chamala et al., 2015), (iii) alternative splice donor site, and (iv) alternative splice acceptor site. In plants, the majority of splice junctions are located in the protein coding sequences (CDS) of genes compared to the UTRs (Marquez et al., 2012). Alternative splicing in these junctions can introduce premature termination codons (PTC), which can change the transcript’s susceptibility to nonsense mediated decay (NMD). It is estimated that NMD is the fate of alternative transcripts for 13% of *Arabidopsis* multi-exon genes (Kalyna et al., 2012). Certain rules have been described that dictate when transcripts may be targeted by NMD, such as an elongated 3’-UTR or the presence of uORFs, but exceptions to these rules exist and it is currently not fully understood in which cases transcripts are effectively degraded (Kalyna et al., 2012). Isoforms that are not degraded by NMD can get translated into proteins with altered (related, distinct, or opposing) functions and/or changes in subcellular localization, stability, enzymatic activity, binding or posttranslational modifications (Carvalho et al., 2013; Kelemen et al., 2013). Alternative splice events that change the transcript’s untranslated regions can impact mRNA stability or translational efficiency (Hughes, 2006; Kalyna et al., 2012).

In humans, it is estimated that 95% of multi-exon genes undergo alternative splicing and that there are on average at least seven alternative splice events per gene (Pan et al., 2008). Human protein-coding genes often have an isoform expressed at a significantly higher level than others and it is therefore referred to as dominant (Trapnell et al., 2010; Gonzalez-Porta et al., 2013). Under certain conditions these genes can change the relative expression levels of their isoforms in events that are defined as Differential Transcript Usage (DTU). Here, we refer to DTU as events in which isoforms gain and/or lose dominance in one condition compared to another. Events that cause the relative abundance order of transcripts to change are called isoform switches and current methods to identify these isoform switches are based on the detection of reversals of relative transcript abundance order between replicated case and control samples (Sebestyen et al., 2015) or in time series (Guo et al., 2017). Here, isoform switches are defined as DTU events where dominance completely shifts from one isoform to another.

For plants, alternative splice events have been found for 40-70% of multi-exon genes in nine angiosperms (Chamala et al., 2015). Isoform dominance, DTU, and isoform switching has also been observed in several plant species. In *Vitis vinifera* (grapevine), 4,069 genes were found to exhibit at least one isoform switch across 124 samples from different tissues, stress treatments, and genotypes (Vitulo et al., 2014). In *Zea mays* (maize), 1,204 and 3,064 genes with isoform switches were observed over the course of development and under drought stress treatment respectively, in a study covering 94 samples (Thatcher et al., 2016). In *Arabidopsis thaliana*, only 812 genes in the TAIR10 annotation (Lamesch et al., 2012) have so far been reported to switch isoforms across 61 samples (Sun et al., 2014). Stress and development have been shown to impact regulation of alternative splicing in *Arabidopsis thaliana* (Filichkin et al., 2010; Reddy et al., 2013; Staiger and Brown, 2013; Filichkin et al., 2015), but the effects on DTU have not yet been studied on a genome-wide scale. It is for example not known what proportion of alternatively spliced genes or which types of genes undergo DTU, in what respect DTU is influenced by treatments or developmental stages, or what the implications are for the protein coding sequences. Furthermore, unknown DTU events can confound gene-level experimental procedures, such as differential expression analyses where an isoform switch can be missed due to an unchanged expression at the gene level, or the creation of overexpression or knockout lines where an unexpected isoform switch can disrupt the function or the expected effect on the production of the gene product. Given the prevalence of alternative splicing in plant genes, there is an increasing need for a dataset or platform to examine the DTU behavior of genes of interest.

Here we present a large scale dataset containing isoform expression levels, isoform dominance calls, and DTU calls for *Arabidopsis thaliana* across 206 public RNA-Seq experiments, covering different combinations of stress and hormone treatments and a wide array of vegetative and reproductive tissues. For this, the AtRTD2 reference transcriptome was used (Zhang et al., 2017), which provides the most extensive and accurate collection of transcripts for *Arabdopsis thaliana* to date and consists of a merge between annotations from TAIR10 (Lamesch et al., 2012), Araport11 (Cheng et al., 2016), AtRTD (Zhang et al., 2015), and a new transcript assembly based on 129 RNA-Seq libraries (Zhang et al., 2017). Based on the generated dataset, we provide a genome-wide overview of DTU in relation to gene function by integrating sample metadata, Gene Ontology annotation, and protein domain analysis. We show that the frequency at which genes undergo DTU and the effect of these events on the protein sequences differ substantially between genes of different evolutionary ages, functions, structure, and expression breath. Finally, we showcase how the detected DTU events can be used to gain further insight in the occurrence and function of alternative splicing for specific genes of interest or to find candidates for functional analysis.

## RESULTS AND DISCUSSION

### Benchmarking the detection of isoform dominance and DTU shows best performance for an Ensemble method

Before quantifying isoform expression levels using public datasets from the Sequence Read Archive (SRA), the detection of isoform dominance and DTU was benchmarked on simulated RNA-Seq datasets. These were generated with the Flux simulator (Griebel et al., 2012), which takes as input the number of RNA molecules (molecule count) to simulate for each isoform and can simulate single-or paired-end reads of different lengths. A compendium of 100 simulated samples was generated, containing molecule counts for all isoforms in the AtRTD2-QUASI *Arabidopsis thaliana* reference transcriptome (Zhang et al., 2017). AtRTD2-QUASI is a modified version of AtRTD2 that has been optimized for isoform expression quantification. Every isoform from genes with two or more annotated isoforms was simulated as dominant in exactly 20 of the samples by assigning it a five times higher molecule count than the non-dominant isoforms on the same gene (see Experimental Procedures). Flux was then used to simulate single- and paired-end RNA-Seq reads of different lengths for each of the 100 virtual samples. The resulting RNA-Seq datasets were processed with three popular expression quantification tools: Kallisto (Bray et al., 2016) and Salmon (Patro et al., 2017), which are alignment-free methods, and Cufflinks (version 2) (Trapnell et al., 2010), which requires reads aligned to the genome as input. Simulated reads were first aligned to the TAIR10 reference genome using STAR (Dobin et al., 2013), which was chosen for its speed and accuracy (Engstrom et al., 2013).

For each simulated RNA-Seq dataset, Cufflinks produced Fragments Per Kilobase of transcript per Million mapped reads (FPKM) values (Mortazavi et al., 2008) whereas Kallisto and Salmon produced Transcripts Per Million (TPM) values (Li et al., 2010). Although the FPKM normalization is optimized for comparison of gene expression within the same sample, whereas the TPM normalization is better suited for comparing expression values between samples, both types of expression values were treated the same (see Experimental Procedures). Isoform dominance was called with each tool individually by setting a threshold on the ratio of an isoform’s FPKM or TPM over the median FPKM or TPM of the gene’s other isoforms in an iterative manner (see Experimental Procedures). This ratio is from here on referred to as the FPK ratio. An Ensemble method was also implemented which calls an isoform as dominant when at least two out of the three individual tools confirm the dominance. DTU was called by first selecting the most common configuration of dominant isoforms for each gene across all simulated samples. Next, any deviation from this configuration (loss or gain of dominance by one or more isoforms) was considered as DTU except when no dominant isoforms were detected (see Experimental Procedures). Finally, a statistical filter step based on Jensen-Shannon distances was used to remove DTU calls if the detected relative abundances of transcripts were too similar to the relative abundances observed in samples with the favorite configuration. It should be noted that alternative and recommended methods exist to detect DTU and isoform switching between two replicated conditions, such as iso-kTSP (Sebestyen et al., 2015) and SwitchSeq (Gonzàlez-Porta and Brazma, 2014), and to detect isoform switching in time series such as TSIS (Guo et al., 2017). These methods were however not applicable in the construction of this compendium of public datasets since it contained a mix of multiple organs and treatments, many of which did not have replicates.

Detection of dominant isoforms was evaluated by calculating recall, precision and F1 scores (the harmonic mean of recall and precision) for each isoform individually by counting the number of correct dominance calls across the 100 simulated samples. The same measures were calculated for each gene individually to evaluate DTU calls. Performance was assessed for each tool separately and for the Ensemble method by plotting the F1 scores in boxplots along with average recall and precision scores (Figure 1). RNA-Seq datasets with different read length (50nt, 75nt, and 100nt), read type (single-end, paired-end), and read numbers (10, 20, 30, 40, and 50 million reads) were examined to test the robustness of the detection method (Figure 1). These ranges of read numbers and read lengths were chosen to reflect the properties of public RNA-Seq datasets (Supplemental Figure S1).

**Figure 1.**
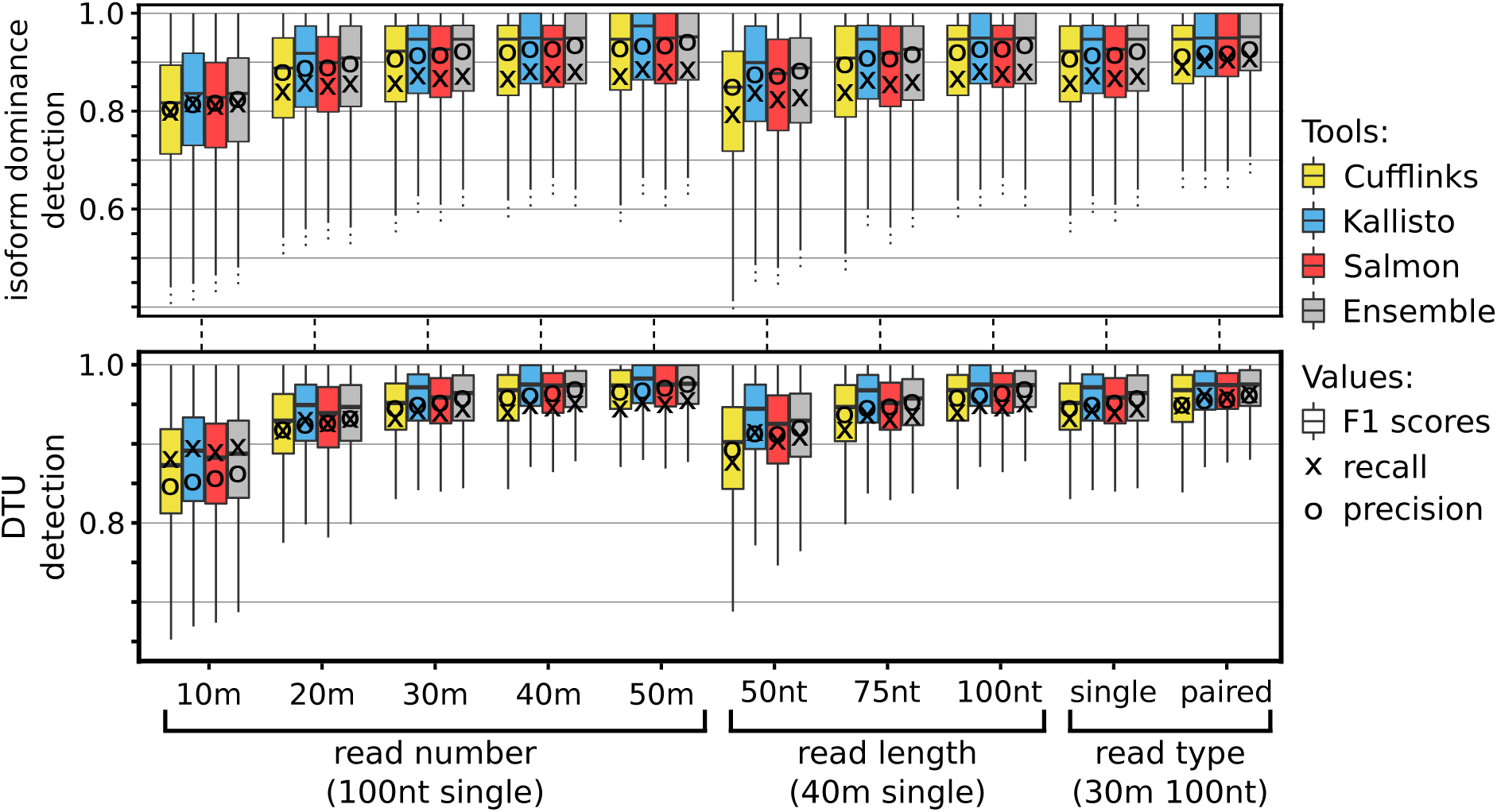
Read properties influence the detection of isoform dominance and DTU. Three properties of reads were examined: the read number (per million), the read length (in nucleotides), and the read type (single-end and paired-end). Boxplots represent F1 scores per isoform for dominant isoform detection and per gene for DTU detection. Average recall and precision are marked by X and O respectively. Boxplot outliers are not shown.

Downsampling the read number showed that performance decreased as the number of reads dropped, but that even with 10 million reads the tools still performed well with the Ensemble method achieving average F1 scores of 80.83% and 86.83% for isoform dominance and DTU detection respectively (Figure 1). Longer reads improved results, likely due to the increased chance for each read to span a discriminative exon-exon junction. Each of the tools showed higher F1 scores for 20 million reads of 100bp compared to 40 million reads of 50bp even though the total number of sequenced nucleotides in both datasets was equal. To test the effect of paired-end sequencing, results for 30 million single-end reads of 100nt were compared to 15 million paired-end read pairs of 2x100nt. The total number of sequenced nucleotides is again equal in both datasets, yet the use of paired-end reads improved average F1 scores of the Ensemble method from 94.47% to 95.67% for DTU detection.

Of the three individual tools, Kallisto achieved the highest precision, recall and F1 scores for each simulated dataset. The ensemble method improved precision and F1 scores of DTU detection by 0.85% and 0.50% respectively across all setups compared to Kallisto and was therefore selected to process the public datasets.

### The number of isoforms per gene and their physical overlap affects dominant isoform detection performance

Over the past few years, the number of known isoforms in the *Arabidopsis thaliana* transcriptome has increased drastically. In November 2010, the TAIR10 annotation contained 27,416 protein coding genes of which 21% had more than one known isoform (on average 2.37 isoforms per gene) (Lamesch et al., 2012). In 2016, two new annotations were released: Araport11 in which 41% of 27,655 protein-coding genes had more than one isoform with on average 2.94 isoforms per gene (Cheng et al., 2016), and AtRTD2 in which 49% of 27,667 protein-coding genes (60% of 22,453 intron-containing) had more than one isoform with on average 4.44 isoforms per gene (Zhang et al., 2017). However, the large number of isoforms per gene in AtRTD2 potentially complicates expression quantification since an increased number of isoforms negatively impacts quantification accuracy (Hayer et al., 2015; Kanitz et al., 2015; Zhang et al., 2017). To investigate if this affected the detection of isoform dominance and DTU, genes were first binned based on their number of transcripts. Next, the average F1 scores of their transcripts (obtained with the Ensemble method) were plotted in Figure 2. As the number of transcripts increased, the F1 scores decreased as expected yet the curve showed two unexpected properties. First, the curve exhibited a stepwise decline where the F1 scores decreased more when incrementing the number of isoforms from an odd to an even number than vice versa. Second, F1 scores appeared better for genes with three isoforms than for genes with only two isoforms, which argues against the assumption that isoform abundance estimation is more complex for genes with more isoforms. The stepwise decline of F1 scores can be explained by the way dominant isoforms were defined. If we consider the calling of dominant isoforms as a classification problem then N isoforms need to be classified in a set of dominant and a set of non-dominant isoforms. Since we stated that no more than half a gene’s isoforms could be simulated as dominant in a single sample, the maximum number of dominant isoforms is N/2. This number only changes when incrementing N from an odd to an even number, which offers an explanation for the uneven decline of F1 scores.

**Figure 2.**
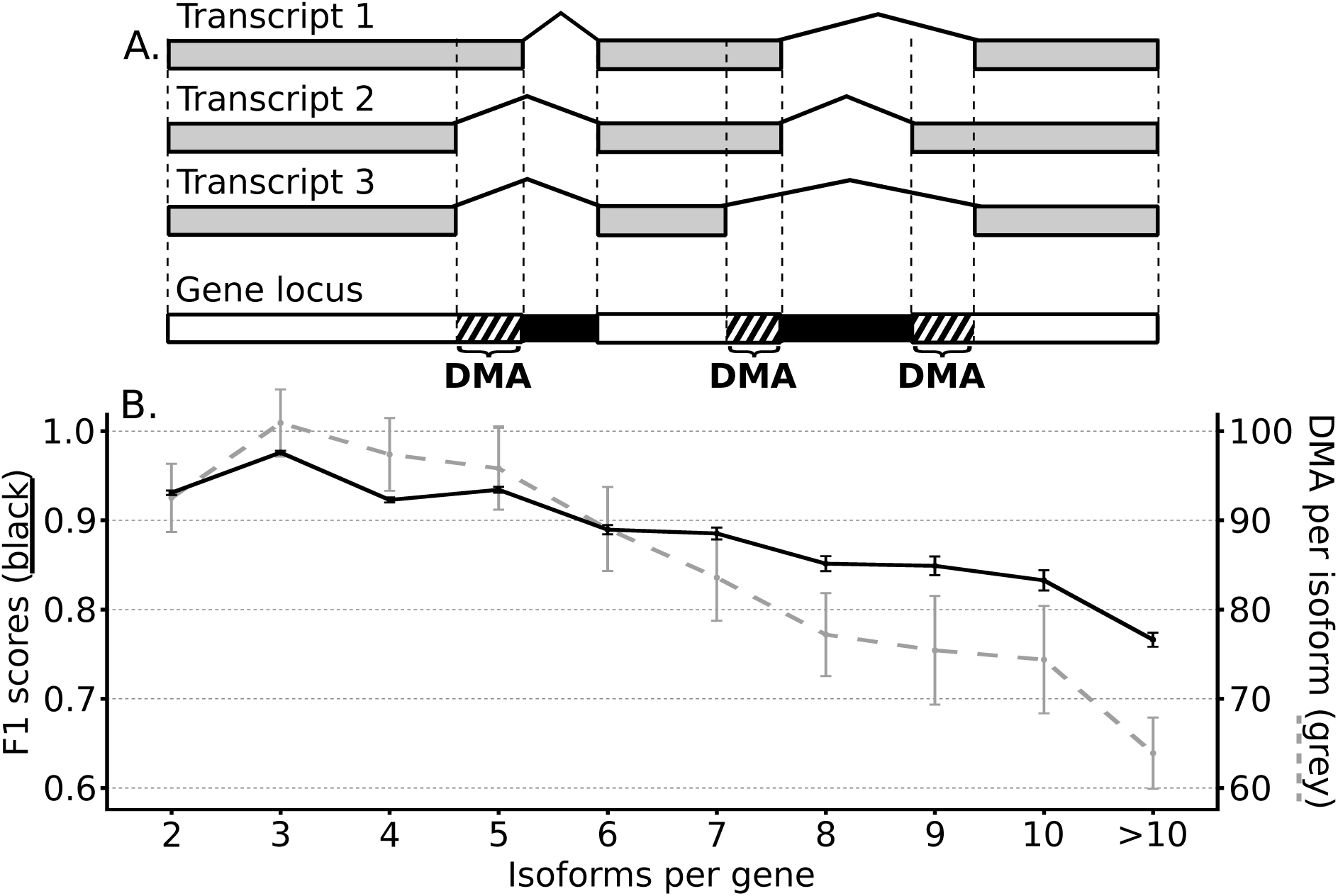
Discriminative Mapping Area length correlates with dominant isoform detection performance. Panel A illustrates the definition of the Discriminative Mapping Area (DMA) of a gene as its combined genomic areas that are only present in a subset of its isoforms. Panel B shows the correlated decline of the DMA length per isoform (grey, in nucleotides) and the average F1 scores of dominant isoform detection (black) for genes with an increasing number of annotated isoforms. DMA per isoform was calculated as the length of a gene’s DMA divided by its number of isoforms. Error flags represent the 95% confidence intervals.

To explain the higher F1 scores for genes with three isoforms compared to genes with only two, we defined the Discriminative Mapping Area (DMA) of a gene as the union of the genomic areas that are only present in a subset of its isoforms (Figure 2). RNA-Seq reads that align to a gene’s DMA can be used by the expression quantification tools to estimate relative abundances between isoforms and so a longer DMA should benefit isoform expression quantification and subsequent dominant isoform detection. To verify this, a Spearman’s rank correlation was calculated between a gene’s DMA per isoform (the DMA length in nucleotides divided by its number of isoforms) and the average F1 score of the dominance detection of the gene’s isoforms (for 30 million single-end reads of 100bp). A significant positive correlation was found (rs=0.26, p<0.001) and this correlation increased to rs=0.35 and rs=0.42 when only genes with fewer than five and four isoforms were considered, respectively. The DMA per isoforms peaks for genes with three isoforms and is lower for genes with two isoforms, which offers an explanation for the difference in F1 scores (Figure 2). Overall, our results indicate that the performance of DTU detection decreases when more isoforms are annotated due to a decrease in DMA length per isoform. However, even for genes with ten isoforms the average F1 score remains high at 83.28%, which demonstrates the robustness of the ensemble method against this decline in DMA lengths.

### Construction and validation of a large-scale isoform dominance and DTU compendium

To generate a large-scale compendium of isoform dominance and DTU calls for *Arabidopsis thaliana*, publicly available RNA-Seq datasets from SRA were first manually curated (see Experimental Procedures). This resulted in a set of 121 samples covering 43 different combinations of treatments and organs (Supplemental Table S1). A recent RNA-Seq library containing 85 samples for 79 organs and developmental stages, as well as time-series for heat, cold, and wounding stresses, was also included (Klepikova et al., 2016). Raw RNA-Seq data for all 206 samples were processed to obtain FPKM and TPM values from Cufflinks, Salmon, and Kallisto (Supplemental Data S1), and isoform dominance and DTU was detected using the Ensemble method (Supplemental Data S2 and S3).

To evaluate if the performance of dominant isoform detection for the public RNASeq datasets was in line with the performance observed for simulated datasets, the isoform dominance calls for the public RNA-Seq samples were compared to a high-resolution reverse transcription polymerase chain reaction (RT-PCR) panel (Simpson et al., 2008), which has previously been used to validate alternative splice events (Marquez et al., 2012; Zhang et al., 2015). This study reports Percent Spliced (PS) values for 34 genes in *Arabidopsis* root, flower, and light- or dark grown seedling. These PS values indicate the relative abundance of transcripts containing either a proximal or distal splice event (Supplemental Table S2). To make these PS values comparable to the categorical output of the dominant isoform detection, the following approach was used: First, to complement the organs examined in the Simpson et al., 2008 study, untreated public RNA-Seq samples were selected for root (14 samples), leaf (37 samples), and flower (20 samples) (Supplemental Table S1). Next, for each gene the number of samples in which isoforms with either the proximal or distal splice event were detected as dominant was counted per organ. These counts were used to calculate a Percent Samples (PSam) values for the distal splice event (see Experimental Procedures). Large or small PSam values indicated that isoforms with the distal or proximal splice event respectively were found to be dominant in most samples. These PSam values were then compared to PS values from Simpson et al., 2008. Large PS values for a distal splice event indicated that this event was dominant in the Simpson et al., 2008 sample, which should be reflected by a large PSam value generated with the dominance calls. Figure 3 shows a scatter plot comparing the RT-PCR PS and the RNA-Seq PSam values for 20 genes that have available PS values for the relevant organs in Simpson et al., 2008 and for which the reported proximal and distal splice events were annotated in AtRTD2. A strong positive correlation was found (Spearman’s rho = 0.80, p <0.001), which confirmed the quality of the isoform dominance calls on public RNA-Seq datasets.

**Figure 3.**
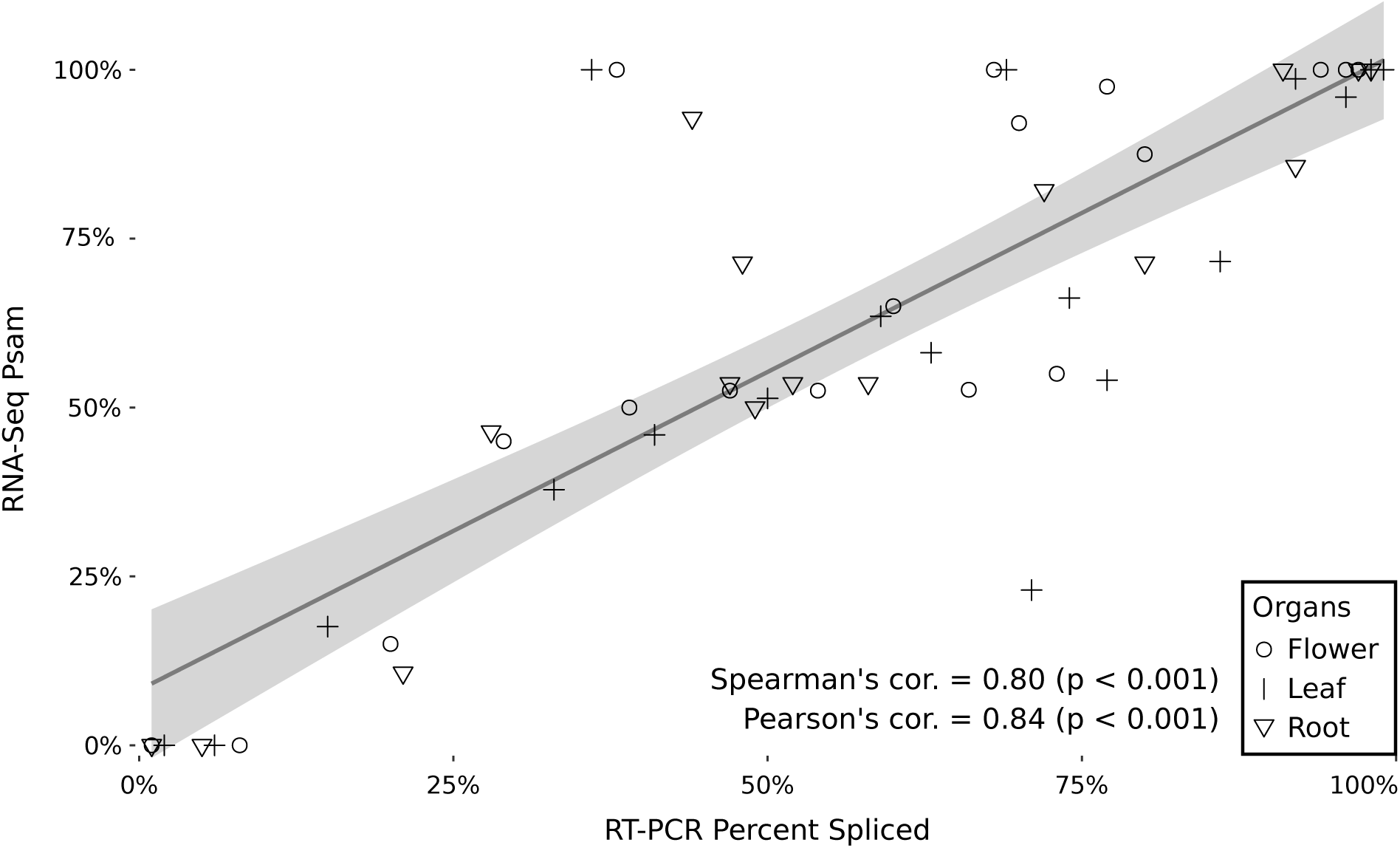
Comparison of isoform dominance calls to RT-PCR Percent Spliced values. RT-PCR Percent Spliced values from Simpson et al., 2008 were compared to Psam values from the dominant isoform detection in flower, leaf, and root samples. Large PSam values indicate that only isoforms containing a specific distal splice event were found to be dominant in most samples, which corresponds to large PS values for this distal splice event. The dark line shows the linear regression between PSam and PS, the grey area indicates the 95% confidence interval. Data shown here are available in Supplemental Table S2.

### Hierarchical clustering of public RNA-Seq samples shows consistent patterns of treatment- and organ-specific DTU

To provide an overview of the samples in the compendium, and to examine the robustness of the detected DTU across replicates, a hierarchical clustering of the samples was performed based on the DTU calls (Figure 4) (see Experimental Procedures). Apart from five exceptions (samples 27, 57, 64, 65, and 108), the remaining 94 replicate samples (connected by red lines in Figure 4) clustered closely together, which demonstrated a high consistency of the DTU calls. High seed dormancy (samples 49-51), drought stress (samples 98-99), and ozone treatment (samples 105-107) resulted in separate clustering of treatment and control samples, which indicates an effect of these treatments on DTU. Samples from five separate studies of biotic stress (*B. cinerea* and *P. syringae* infection) on leaves clustered in two groups (samples 43-48 and 55-59), which indicated reproducible DTU in response to biotic stress as well. Leaf samples from the Klepikova et al., 2016 study treated with heat, wound, and cold stress also separated in three clusters (samples 144-148, 149-153, and 156-159 respectively). Most samples from the Klepikova et al., 2016 study formed one large group, separate from other samples from similar tissues and treatments, suggesting the presence of a laboratory bias in the observed DTU events. Within the Klepikova et al., 2016 study however, most samples from similar organs clustered together. Samples 160-168 formed a rather heterogeneous cluster of reproductive and vegetative organs and organ parts, yet these samples were all taken from mature (samples 160-163) and senescent (samples 164-168) organs, which indicates a role of DTU in aging and senescence. Overall, the tight clustering of replicates and the separation of samples from different organs and treatments revealed the presence of reproducible DTU patterns related to development and in response to e

**Figure 4.**
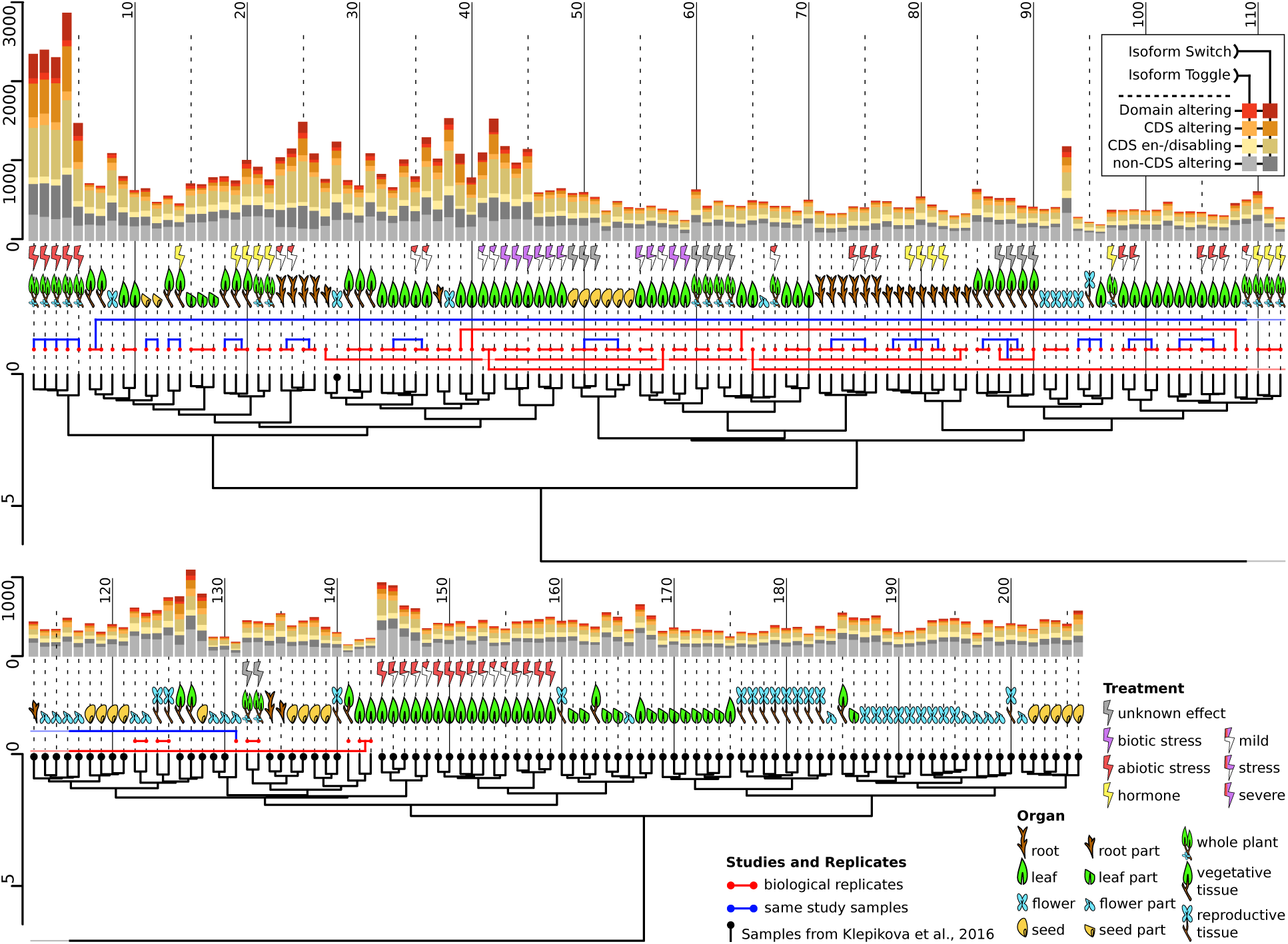
Hierarchical clustering of public RNA-Seq samples based on detected DTU. 206 Public RNA-Seq samples from the Sequence Read Archive were clustered based on DTU calls generated with the Ensemble method. Biological replicates are connected by red lines, samples from the same study are connected by blue lines. Samples marked by a black dot are from a single large-scale RNA-Seq study by Klepikova et al., 2016. Sample metadata regarding organ types and treatments are presented as illustrations. For each sample, the number of genes with a detected DTU and the type of the observed DTU are shown in stacked barcharts. Detailed metadata for each sample can be found in Supplemental Table S1.

### DTU occurs in genes across a wide functional landscape and frequently alter protein sequences and domains

The bar charts in Figure 4 represent the number of genes undergoing DTU in each sample. DTU events where at least one isoform lost dominance while at least one other isoform gained dominance were classified as isoform switches (Figure 5) and are represented by darker shades in the bar charts.

**Figure 5.**
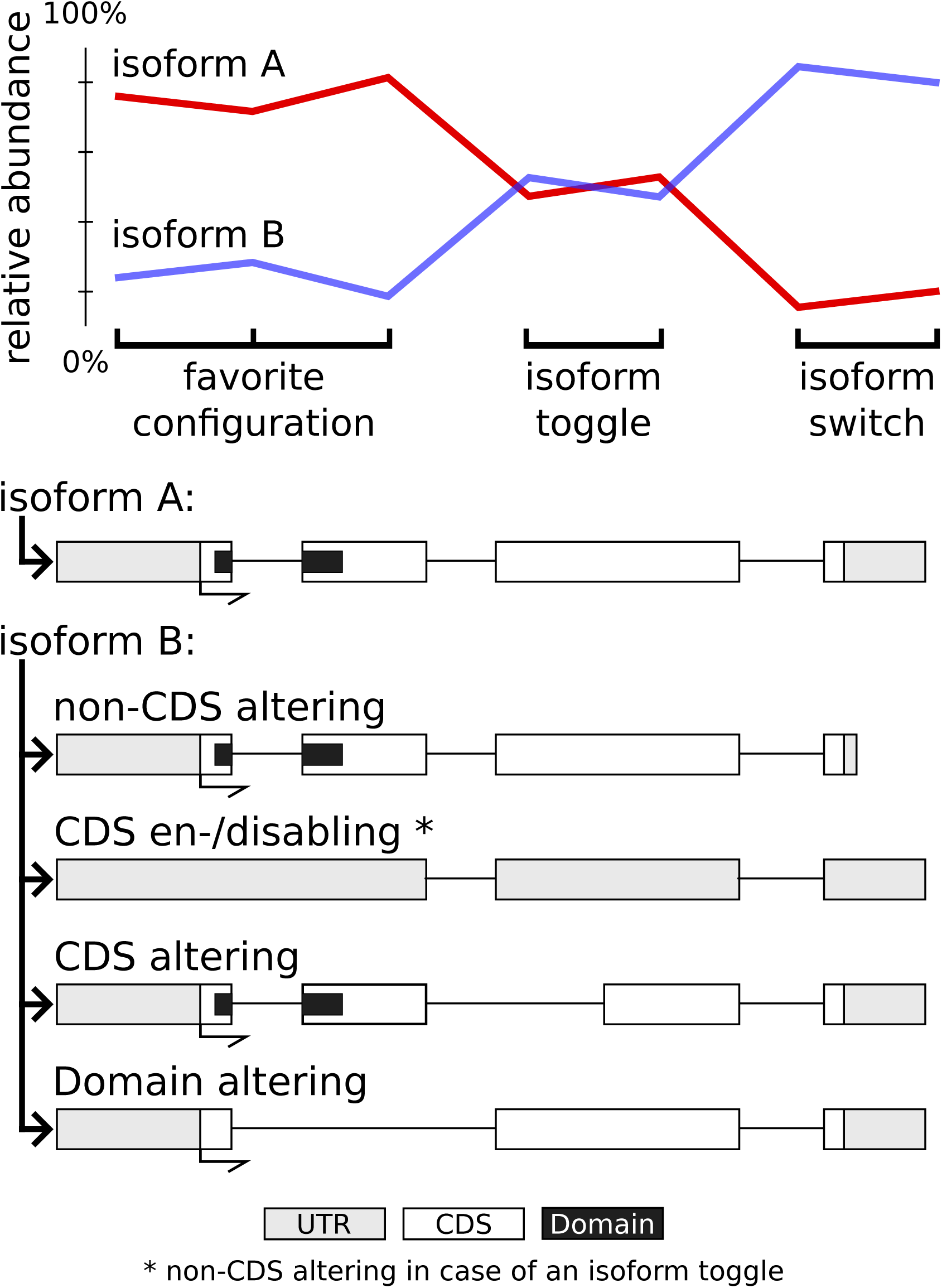
Classification of DTU events. The line plots show the relative abundances of two hypothetical isoforms for a gene undergoing DTU events with and without an isoform switch. The cartoons show the structure of isoform A and possible structures of isoform B. Each structure of isoform B represents a different class of DTU event.

All DTU events were further divided into four classes based on their effect on the protein coding sequences (CDSs) and the protein domain contents of the final gene products (Figure 5) (see Experimental Procedures): (i) events that did not affect the CDSs (non-CDS altering), (ii) changes from or to a configuration where none of the dominant isoforms were annotated with a valid CDS (CDS enabling/disabling), (iii) events that altered CDSs without changing their protein domain content (CDS altering), and (iv) events that altered the CDSs and also changed their protein domain content (domain altering). The CDS data were obtained from AtRTD2 and are the result of an algorithm that fixes the translation start site AUG at the same position for each isoform rather than searching for the longest open reading frames (Zhang et al., 2017). This avoids the erroneous assumption that translation can start at a downstream AUG if the authentic one leads to a PTC (Brown et al., 2015). Isoforms with a CDS shorter than 300 nucleotides were not considered to be protein coding since these transcripts are likely targeted by NMD (Kalyna et al., 2012). It should be noted that this threshold of 300 nucleotides does not eliminate all possible NMD targets since longer transcripts can be susceptible to NMD as a result from other characteristics, such as elongated 3’-UTRs, the presence of certain introns in the 3’-UTR or 5’UTR, or upstream open reading frames (Kalyna et al., 2012). Isoforms that were the result of an intron retention event were also not considered to be protein coding since this type of splice event can cause transcripts to be retained inside the nucleus, which prevents translation (Gohring et al., 2014; Brown et al., 2015).

In total for the 14,916 genes with more than one isoform in AtRTD2-QUASI, 8,148 were found to undergo DTU events. Isoform switches were observed for 7,423 of these genes in at least one of the samples relative to the favorite configuration. For the remaining 725 genes, none of the DTU events involved an isoform switch. Of all observed DTU events, 46.9% were isoform switches.

On average per sample, 587 genes underwent DTU, which resulted in an altered CDS and domain in 15% and 7% of the cases respectively. In 31% of the cases, a CDS enabling/disabling event was observed. These can result from a change in dominance of isoforms with unknown CDSs, a premature termination codon which makes them potential targets of NMD, or an intron retention event. Of all observed DTU events, 48% involved the gain or loss of dominance of at least one isoform with an intron retention event, the most common type of alternative splice events (Chamala et al., 2015). The largest number of DTU events were found in samples 1-5, which are samples of whole seedlings placed under severe abiotic stresses to study alternative splicing in *Arabidopsis* (Filichkin et al., 2010). The full data set of DTU events together with detailed sample metadata is available in Supplemental Data S3 and Supplemental Table S1.

### Genes with frequent, rare, or no DTU differ in evolutionary ages and functions

Counting how frequently each gene underwent DTU revealed that the majority of genes change isoform dominance at least once. Out of the 14,916 genes with two or more isoforms in AtRTD2-QUASI, excluding 1,331 genes with expression in less than five samples, 5,490 genes did not show DTU in any of the 206 samples (no DTU). For the remaining genes, DTU frequencies were calculated as the percentage of expressed samples in which a DTU is observed (Figure 6). These genes were then grouped into rare DTU and frequent DTU genes by setting the median DTU frequency of 5.27% as a threshold. To examine if the occurrence of DTU can be linked to gene functions or to the evolutionary age of the genes, gene set enrichment analyses were performed on the non-, rare-, and frequent DTU genes (Figure 6) (Supplemental Table S3). For Gene Ontology (GO) terms, enrichments were calculated for each term in the GO slim annotation, including annotations for all evidence types (Gene Ontology, 2015). For the evolutionary gene age analysis, each gene was assigned to one of five phylostrata: Viridiplantae, Embryophyta, Angiosperms, Eudicots, and Brassicaceae. Phylostrata describe for each gene the lowest common ancestor of the species that contain a homolog of the gene (Proost et al., 2015) (see Experimental Procedures).

**Figure 6.**
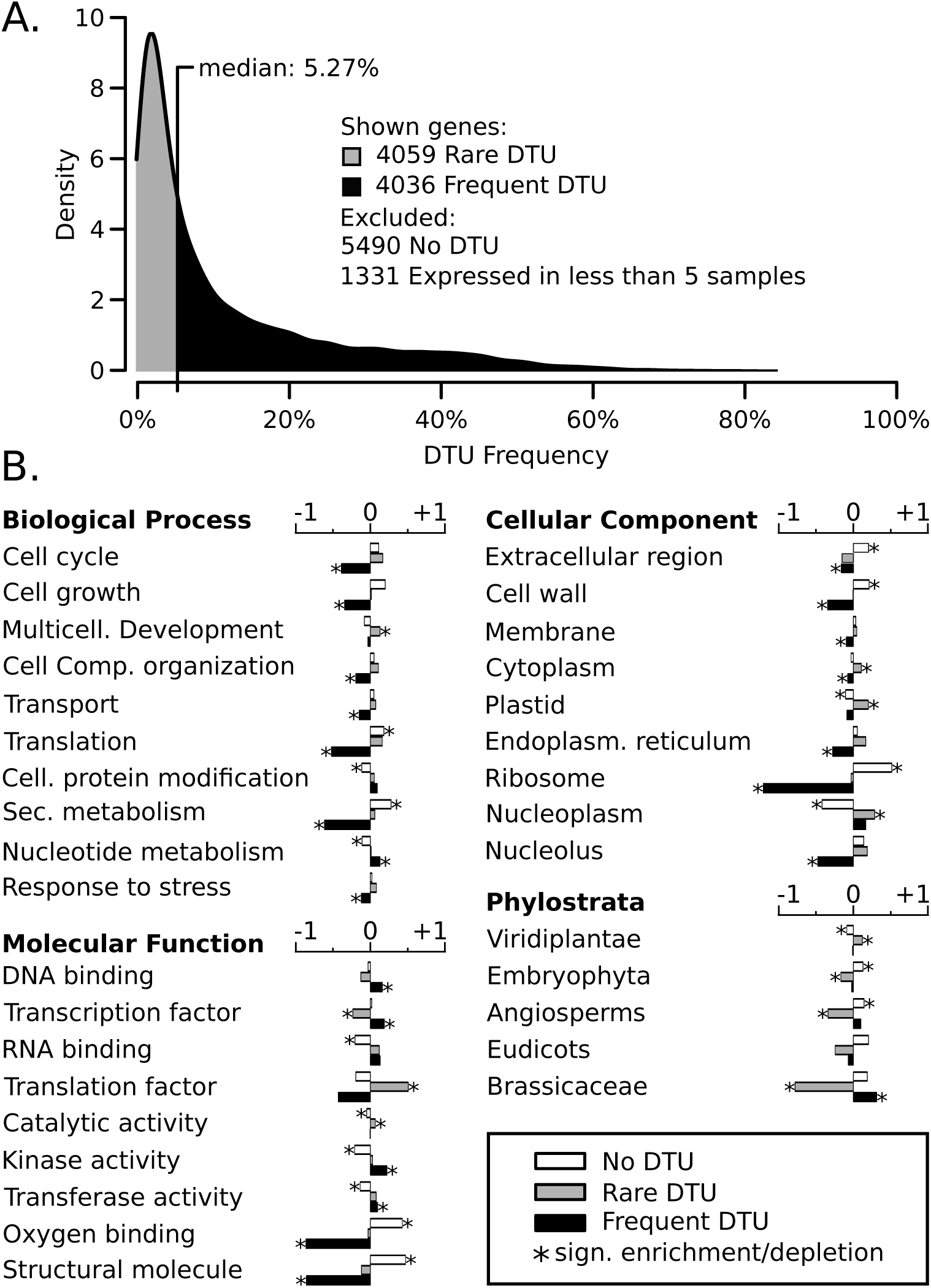
Gene set enrichment analyses of genes with different DTU frequencies. For each gene the DTU frequency was calculated as the ratio of samples in which the gene undergoes DTU over samples in which the gene is expressed. Panel A shows a density plot of DTU frequencies of genes with at least one DTU event. Panel B shows log2 enrichment folds for a selection of Gene Ontology terms and gene phylostrata for genes with different DTU frequencies. Negative values indicate depletions. Asterisks indicate enrichments or depletions that were significant according to a hypergeometric test with FDR controlled at 0.05. Full enrichment results can be found in Supplemental Table S3.

A selection of GO terms that were significantly enriched (based on a hypergeometric test with Benjamini-Hochberg correction at FDR=0.05, see Experimental Procedures) are shown in panel B of Figure 6. Genes without DTU were most noticeably enriched for translation, secondary metabolism, ribosomal and cell wall components, oxygen binding, and structural molecules. All of these terms were strongly depleted in the frequent DTU genes. Rare DTU genes showed enrichment for translation factor activity, multicellular development, and plastid and nucleoplasm localized genes. Frequent DTU genes were enriched for nucleotide metabolism and transcription factor, transferase, and kinase activity. Overall, these results suggest that genes without DTU are more often involved in housekeeping functions, whereas genes that frequently undergo DTU more often take on regulatory functions. Enrichment analyses using phylostrata information showed that the oldest genes, conserved within Viridiplantae, are depleted for no DTU and enriched for rare DTU genes, whereas younger genes showed the opposite trend. Frequent DTU genes were found to be enriched for young Brassicaceae-specific genes, whereas rare DTU genes were depleted.

### CDS altering DTU occurs disproportionately across genes with varying properties

To examine if genes with different characteristics exhibit different DTU behavior, genes were first grouped in five categories: (i) genes that underwent no DTU, and genes that underwent at least one (ii) non-CDS altering, (iii) CDS toggling, (iv) CDS altering, or (v) domain altering DTU event (Figure 5). Each gene was only placed in the most specific applicable category (e.g. genes with both CDS toggling and domain altering DTU events were placed only in the domain altering DTU event category). Next, genes were binned based on a series of properties: (i) the frequency at which they underwent DTU, (ii) the number of annotated isoforms, (iii) the number of exons, (iv) the expression breadth as the number of samples the gene was expressed in,(v) the number of Protein-Protein Interaction (PPI) partners, (vi) the phylostratum of the gene, and (vii) the number of paralogs the gene has in *Arabidopsis thaliana* (see Experimental Procedures) (Figure 7).

**Figure 7.**
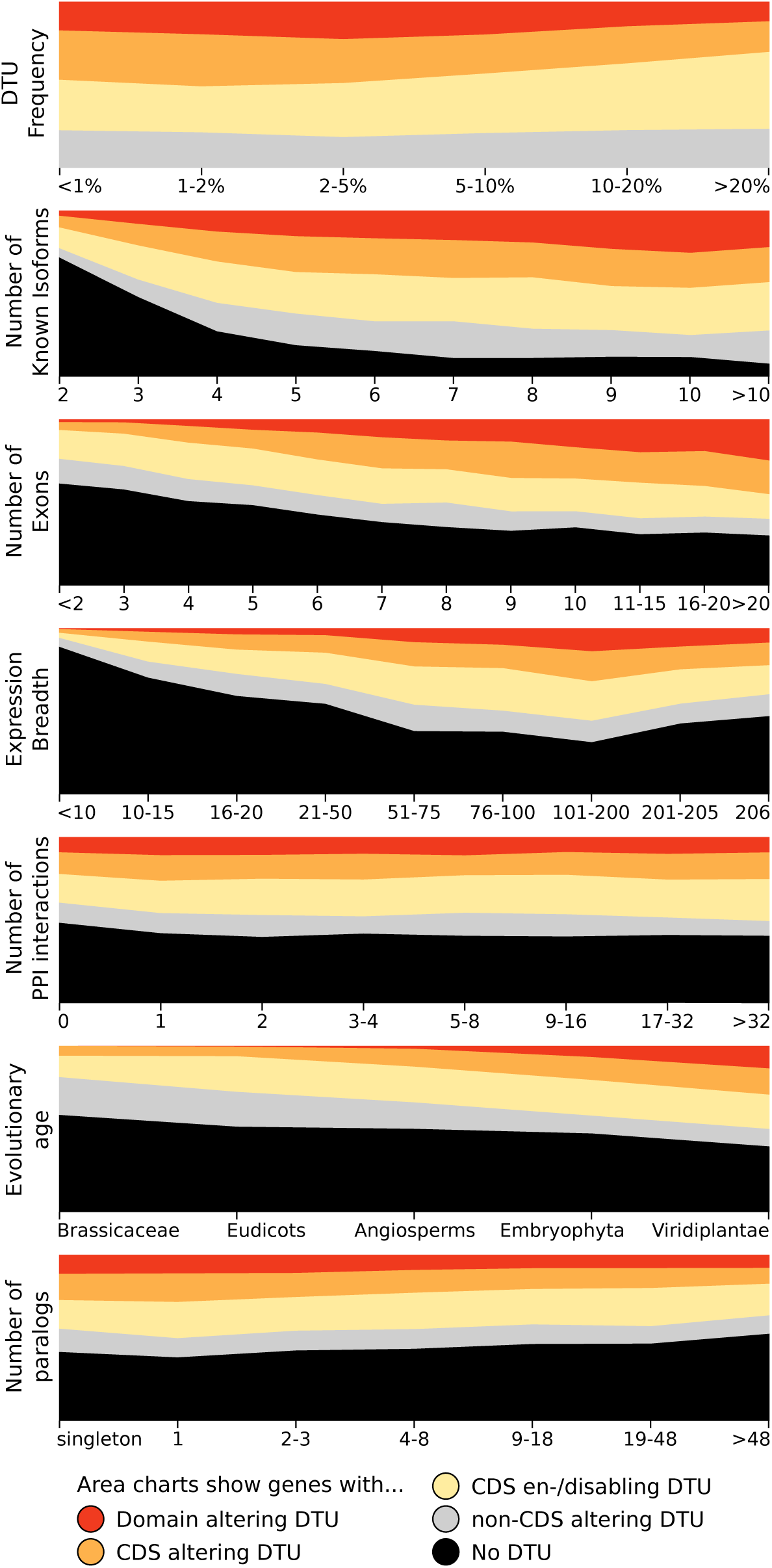
Gene properties influence types of DTU. Genes were binned based on a series of properties: (i) the frequency at which they undergo DTU, (ii) the number of annotated isoforms, (iii) the number of exons, (iv) the expression breadth as the number of samples the gene was expressed in, (v) the number of Protein-Protein Interaction (PPI) partners, (vi) the evolutionary age of the gene, and (vii) the number of paralogs the gene has in *Arabidopsis thaliana*.

Genes with frequent DTU were mostly found to undergo CDS enabling/disabling DTU, whereas rare DTU genes more often showed CDS- and domain altering DTU, indicating that genes that alter their coding sequences tend to do so sparingly (Figure 7). When comparing genes with a different number of annotated isoforms, genes with only two isoforms were less often found to undergo DTU compared to genes with more than two isoforms. Similarly, genes with more exons were more often found to undergo DTU, and a larger portion of these genes showed domain altering DTU. Possibly, both a larger number of isoforms and exons per gene simply increases the number of opportunities for an alternative splice site to occur, which could explain these trends. For the expression breadth it was found that genes with expression in most but not all samples showed the highest proportion of genes with DTU. Reversely, genes with more specific expression in less than 20 samples showed less DTU. Genes expressed throughout the compendium also showed less DTU, which concurred with the previous observation that genes with housekeeping functions, which are typically expressed constitutively, were enriched in genes without DTU events. A strong difference in the categories of DTU was found for genes of different phylostrata. The younger Brassicaceae- and Eudicot-specific genes showed less CDS- or domain altering DTU compared to older genes. The largest number of CDS and domain altering DTU was found for the oldest genes, conserved throughout Viridiplantae. Previously, it has been shown that alternative splicing events are mostly lineage-specific (Chamala et al., 2015; Zhang et al., 2015). Our results suggest that in younger, lineage-specific genes alternative splicing is mostly used as a regulatory mechanism to modulate the level of protein production, whereas in older genes it more often directly affects the functions of the proteins by altering their sequences. Future work will be needed to determine if these alterations to the coding sequences involve these conserved alternative splice events, or if each plant species can produce their own set of new proteins out of old genes. Finally, DTU was compared between genes with a different number of paralogs to study the relationship between gene duplication and DTU. The relationship between gene duplication and alternative splicing has been studied extensively, but is not fully understood (Iniguez and Hernandez, 2017). Singleton genes in rice were found to produce less alternatively spliced isoforms than genes with paralogs (Lin et al., 2008) but for human genes, the reverse trend was observed (Roux and Robinson-Rechavi, 2011). Our findings show that in *Arabidopsis thaliana*, genes with more paralogs are slightly less likely to undergo DTU compared to genes with no or fewer paralogs.

### Case studies showcase the potential of DTU detection for gene function analysis

To demonstrate how the DTU dataset can be used to gain insight in gene function and regulation, the compendium was searched for genes with DTU events that were specific for certain organs or treatments. Three genes with interesting DTU patterns were selected for further examination (Figure 8).

**Figure 8.**
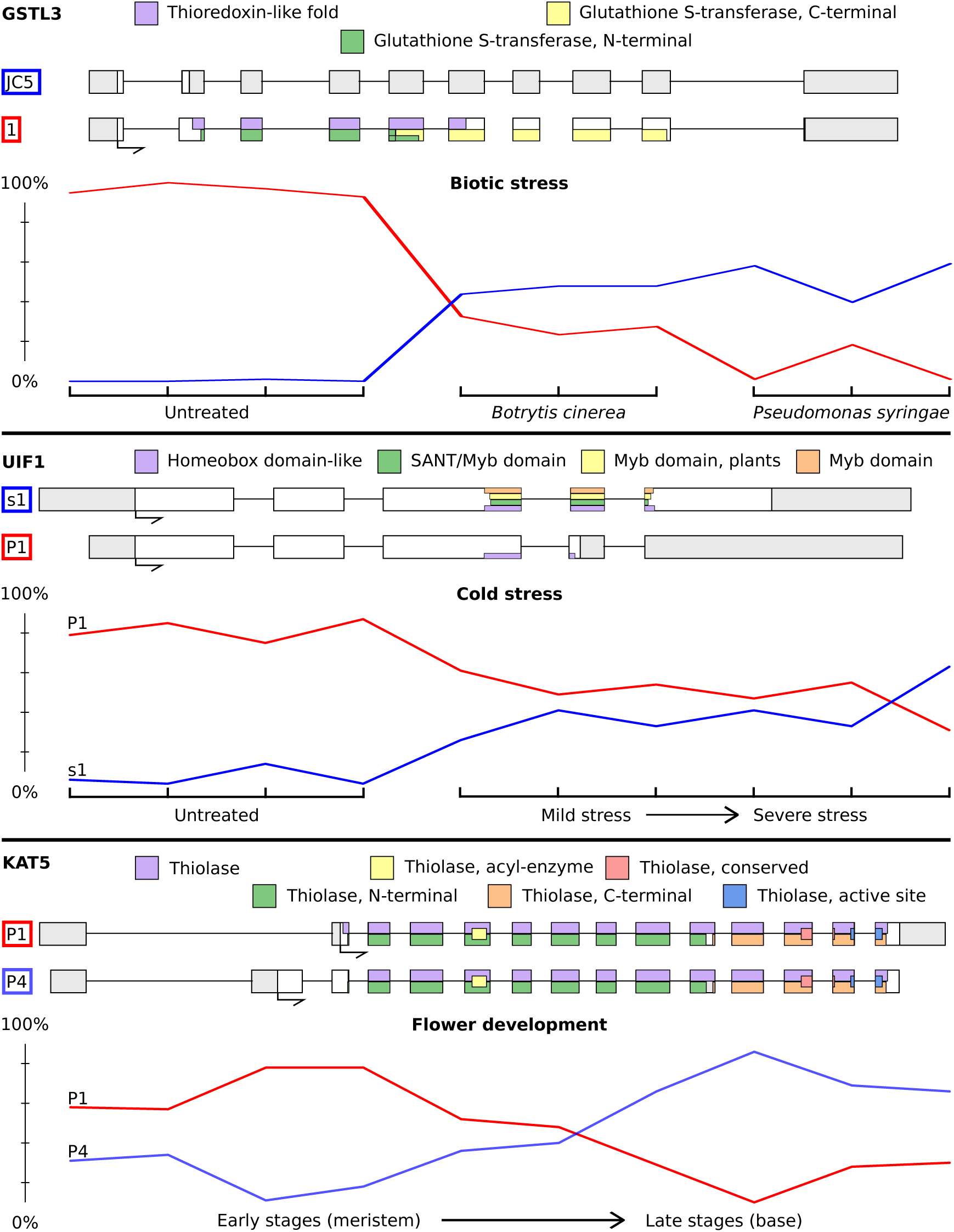
Examples of genes with condition-specific DTU. For each gene, the structure of the involved isoforms and their encoded protein domains are shown. UTR and CDS are shown as grey and white areas respectively. Line plots represent expression levels of the isoforms as a percentage of the total gene expression (y-axis, TPM values from Kallisto were used) across a selection of samples from the public dataset (x-axis). The samples used for each gene are indicated in Supplemental Table S1.

The first gene was GSTL3 (AT5G02790), a member of the lambda glutathione transferases, which can modify cysteinyl residues of proteins in response to chemical stress to directly regulate their activity (Dixon et al., 2005). GSTL3 was expressed in almost all of the 206 public samples and the AT5G02790.1 isoform was found to be dominant in all but six samples from two independent studies: three *Pseudomonas syringae* infected and three *Botrytis cinerea* infected leaves. The overall gene-level expression of GSTL3 does not change in these biotic stress samples, yet in each of them an isoform switch was observed where dominance switched to AT5G02790_JC5 (Figure 8.A). In this isoform, an alternative splice acceptor site introduced a premature termination codon which shortened the reading frame to 13 amino acids, likely causing the isoform to be degraded by NMD. GSTL3 is currently not known to be involved in pathogen response, yet the highly specific nature of this isoform switch would make it and its targets excellent candidates for further study.

A second gene that was examined was *UIF1* (*AT4G37180*), which is a member of the GARP2 G2-like subfamily of transcription factors and is known to control floral meristem activities (Moreau et al., 2016). The *AT4G37180_P1* isoform, which contains a DNA-binding Myb domain, was found to be dominant in most samples. In cold stress samples from two independent studies however, a DTU event was detected where the *AT4G37180_s1* isoform gained dominance (Figure 8.B). This isoform has an elongated fourth exon which shifts the reading frame and introduces a stop codon. This results in a shorter coding sequence with a truncated Myb domain (Figure 8.B). A possible consequence of this is that the protein can still bind its interaction partners but will no longer be able to initialize transcription and might thus act as an inhibitor. An analogous process has previously been described for the starch metabolism regulating transcription factor *IDD14* (Seo et al., 2011). *UIF1* has currently not been annotated with any GO terms that suggest a role in stress response, though given its involvement in floral meristem activities, the detected DTU event makes it an interesting candidate for further functional analysis regarding its role in cold response.

A third gene with interesting DTU events was *KAT5* (*AT5G48880*). It encodes a 3-ketoacyl-CoA thiolase with known involvement in inflorescence meristem development (Wiszniewski et al., 2014). The gene was previously known to undergo alternative splicing, resulting in two versions of the *KAT5* protein: KAT5.2 which contains a type 2 peroxisomal targeting sequence and is located in the peroxisomes, and KAT5.1 which lacks the targeting sequence due to an exon skip event and is located in the cytosol (Carrie et al., 2007). Published analyses with KAT5.1- and KAT5.2-GUS lines reported that KAT5.1 was active from early stages of flower development but was absent beyond stage 12 of development, whereas KAT5.2 activity was observed during the middle stages (9-12) (Wiszniewski et al., 2012). DTU detection on the flower development time series samples from the Klepikova et al., 2016 study confirmed this pattern as a clear switch was observed from *AT5G48880_P1* (KAT5.1) in young flowers at the meristem to *AT5G48880_P4* (KAT5.2) in older flowers near the base of the plant (Figure 8.C). Overall, these examples showcase that the DTU calls generated in this study can be applied to gain both confirmatory results and further insights in the function and regulation of alternative splicing in *Arabidopsis thaliana*.

## EXPERIMENTAL PROCEDURES

### Simulation of RNA-Seq data

The compendium of 100 virtual samples was generated by assigning molecule counts to each transcript in the AtRTD2-QUASI reference transcriptome of *Arabidopsis thaliana* (Zhang et al., 2017). This was done for each sample individually as follows: first, a molecule count was assigned to each gene by randomly sampling from estimated molecule counts. These estimates were calculated by scaling gene-level raw read counts so that the total sum of molecule counts equaled 5 million. Raw read counts were obtained by processing RNA-Seq data from a random experiment on the Sequence Read Archive (SRX1796284) with Salmon (see Isoform expression quantification). Secondly, the molecule counts were increased by 50 for each gene and multiplied by the number of known transcript isoforms for the gene. This ensured that all transcripts were expressed in the simulation. Thirdly, molecule counts for each gene were distributed among its transcripts so that non-dominant isoforms had an equal number of molecules and dominant isoforms had five times more. Finally, the molecule counts were scaled again so that the total sum of molecules amounted to 5 million per sample. Which isoforms were dominant in which samples was randomly selected so that each transcript was dominant in exactly 20 of the 100 samples and less than half a genes’ transcripts were dominant in the same sample.

The Flux simulator (v1.2.1, Flux Library 1.22) (Griebel et al., 2012) was used to simulate FASTQ files with RNA-Seq reads of different type (single-end, paired-end) and length (50nt, 75nt, 100nt) for each of the 100 virtual samples. The default error model for reads of 76 nucleotides was used to simulate sequencing errors. Poly-A tail simulation was disabled as this caused bad performance of expression quantification for paired-end reads. Other parameters were set to default. 60 Million reads were simulated and aligned with STAR (see Isoform expression quantification) for every combination of read type and length. Datasets of 50, 40, 30, 20, and 10 million reads were obtained by random downsampling from the aligned reads. Only primary alignments were kept for the simulated reads.

### Selection of public RNA-Seq datasets

The Sequence Read Archive (SRA) (Kodama et al., 2012) was queried on October 1^st^ 2015 to obtain all studies which had an experiment that matched following criteria: LIBRARY_STRATEGY="RNA-Seq", LIBRARY_SOURCE="TRANSCRIPTOMIC", and STUDY_TYPE="Transcriptome Analysis". This resulted in 214 studies comprising 2799 experiments. These were manually curated to select samples suited for dominant isoform detection: only experiments on Col-0 plants were considered, experiments performed on mutants or transgenic plants were excluded. Samples with fewer than 15 million reads were discarded. Only experiments with Illumina reads were used. Experiments performed on the samples with the same accession on SRA were considered to be technical replicates and their reads were merged to obtain a single set of reads per sample. These steps resulted in a final set of 121 samples for 43 different combinations of treatments and plant tissues (Supplemental Table S1). A recent dataset, containing 85 samples from different organs, developmental stages, and stress treatments was added as well (Klepikova et al., 2016). Samples from the Klepikova et al., 2016 study were available as replicates of two experiments, each performed on separate pools of 15 individual plants, uploaded under the same sample accession to SRA. These replicates were also merged in order to reduce redundancy in the compendium. This brought the total number of samples in the compendium to 206.

### Isoform expression quantification

Reads were aligned to the TAIR10 *Arabidopsis thaliana* reference genome with STAR (v2.4.0j) (Dobin et al., 2013). Expression quantification was performed with Cufflinks (v2.2.1) (Trapnell et al., 2010), Kallisto (v0.43.0) (Bray et al., 2016), and Salmon (v0.6.1) (Patro et al., 2017). For STAR, reads aligning to non-canonical splice junctions not present in known transcripts were removed. The maximum number of multi-mappings per read was set to 15 and reads were allowed to align with 10% mismatches. The maximum overhang on each side of a splice junction was set to 100 and the minimum and maximum intron lengths were set to 5 and 6000 respectively. BAM files needed by Cufflinks were obtained by sorting and converting the SAM files with SAMtools (v1.1) (Li et al., 2009). For the simulations, FASTQ files needed by Kallisto and Salmon were created directly from the SAM file with a custom script. This ensured that all three tools were given the same reads as input.

To process the public RNA-Seq data, SRA files were downloaded from SRA (Kodama et al., 2012) and converted to the FASTQ format using fastq-dump from the SRA toolkit (v2.4.4). FastQC (v0.11.2) (Andrews, 2010) was used to detect overrepresented adapter sequences, which were subsequently clipped with fastx_clipper from the FASTX toolkit (v0.0.13) (Gordon, 2009). Reads shorter than 20 nucleotides after adapter clipping were discarded. The resulting FASTQ files were used as input for Kallisto and Salmon directly, and not aligned first as was done for the simulated data.

Cufflinks, Kallisto, and Salmon were run with default parameters. The index for Kallisto and Salmon was built with a k-mer length of 19 nucleotides. The libtype option for Salmon was set to “"U" and "UI" for single-end and paired-end reads respectively. For Kallisto, the estimated average and standard deviation of the fragment lengths were set to 200 and 20 respectively. Detection of novel transcripts was disabled for Cufflinks.

### Detection of dominant isoforms

Dominant isoforms were detected by calculating the ratio of each isoform’s expression value (FPKM or TPM) over the median expression values of other isoforms from the same gene and calling an isoform as dominant if this FPK ratio exceeded a threshold of 2.2. This threshold was chosen for its optimal F1 scores in a separate benchmark (Supplemental Figure S2). By calculating ratios of FPKM and TPM values, the “per million” scaling factor that differentiates these two normalizations is cancelled out, rendering the ratios from FPKM and TPM values equivalent to calculating the ratio of Fragments Per Kilobase of transcript (FPK) values. Alternative versions to calculate the ratios were tested in a separate benchmark, which also included a method that places a cutoff on an isoforms Percentage of Gene Expression (PGE) value to determine if it is dominant (Supplemental Figure S2). Here, an isoform’s PGE value was calculated as the isoform’s expression value over the sum of expression values of all the gene’s isoforms. One disadvantage of this PGE method is that a custom threshold is required for each number of isoforms. Another disadvantage of this method is that two isoforms with similar expression levels would get a different dominance call if the threshold happens to fall in between them. This situation is less likely for the ratio method since the value of the threshold there is dictated by the other isoforms and would require a very specific configuration of relative abundances to occur. In an attempt to solve both issues for the PGE method, an exhaustive optimization of PGE cutoffs for each number of isoforms was performed and a version of the PGE method was tested that removed dominance calls for isoforms with a PGE value closer than 0.05 to the PGE value of a non-dominant isoform. No version of the PGE method was however able to outperform the ratio method (Supplemental Figure S2).

For the ratio-based method, a ratio of 2.2 was chosen as a threshold for detection based on benchmarks (Supplemental Figure S2). Setting this threshold on the FPK ratio only once would however result in false positives for genes with multiple unexpressed isoforms. If for example a gene with four isoforms has two lowly expressed isoforms with an expression value of 1 (*U1* and *U2*), one moderately expressed isoform with an expression value of 20 (*M*), and one highly expressed isoform with an expression value of 500 (*H*), then *M* would falsely be called as dominant along with *H*. This results from using the median expression of *U1*, *U2*, and *H* to calculate the FPK ratio of *M* which would be equal to 20/median(1, 1, 500) = 20, which exceeds the threshold of 2.2. To avoid these situations, an iterative approach was used where the FPK ratios are recalculated and re-evaluated after removing non-dominant isoforms. In the example of the four isoforms this would result in the removal of *U1* and *U2* after the first iteration and the recalculation of the FPK ratio of *M* as 20/median(500)=0.04 in the second iteration, which no longer exceeds the threshold. If no dominance can be found in the first iteration, all isoforms are classified as non-dominant. If no dominance can be found in a later iteration, the isoforms that were found to be dominant in the previous iteration are returned as the final dominant isoforms. Transcripts with expression values lower than 5.0 are classified as non-dominant by default. A gene was said to be expressed when any of its transcripts was given an expression value greater than or equal to 5.0 by at least two of the three quantification tools.

### Comparison to RT-PCR

In order to compare RT-PCR PS values of certain splice events from Simpson et al., 2008 to detected dominance of transcripts, PSam values were calculated. For this, the number of samples from a specific organ were selected where only isoforms with the distal splice event were dominant (d), only isoforms with the proximal splice event were dominant (p), and where no isoforms were dominant or both isoforms with the distal and proximal splice event were dominant (b). The PSam for a gene was then calculated as (d+(b/2))/(d+p+b). Large PSam values indicate that isoforms with the distal splice event were detected as dominant in most samples, which was expected to be reflected by large PS values of the distal splice event in the RT-PCR panel. Sample counts, PSam, and PS values for all 20 genes shown in Figure 3 are available in Supplemental Table S2.

### Detection and classification of DTU

Based on the transcript-level dominance calls, DTU events were detected for each gene with more than one transcript isoform. For this, an ‘isoform dominance configuration’ was defined as the set of isoforms that are detected as dominant for a gene in a given sample. The ‘favorite configuration’ of each gene was then defined as the most frequently occurring configuration in all 206 public samples (or the 100 simulated samples in the simulations). Any deviation from this configuration, by one or more isoforms losing or gaining dominance, were considered for DTU, except when no dominant isoforms were found. Next, a statistical test was applied to compare relative abundances of isoforms in potential DTU events to the relative abundances observed in the favorite configurations. For this, the Jensen-Shannon distances between relative abundances in the favorite configurations were used to calculate an average distance and a 99% confidence interval. Only if the average Jensen-Shannon distance between the relative abundances observed in a potential DTU event and the relative abundances of samples in the favorite configuration exceeded the 99% confidence interval, the DTU event was called. This test filtered out cases where small differences in expression caused a change in detected dominance by crossing the detection threshold.

If in a detected DTU event at least one isoform lost dominance and at least one other isoform gained dominance compared to the favorite configuration, the DTU event was called an isoform switch since the gene switches expression from one isoform to another. In the other cases, a set of isoforms either gained or lost dominance, which means that the gene merely toggled expression on or off for these isoforms (Figure 5).

All DTU events were further classified based on their effect on the protein coding sequences (CDSs) and protein domain contents of the isoforms in the new configuration compared to the favorite configuration (Figure 5). For this, the set of unique CDSs of the dominant isoforms in each configuration was determined. Similarly the set of unique protein domain combinations in these coding sequences was extracted. When DTU occurred and the new configuration had a changed set of CDSs or protein domain combinations compared to the favorite configuration, this event was classified as ‘CDS altering’ or ‘domain altering’ respectively. Since a loss or gain of dominance for transcripts without a valid CDS does not change the set of CDSs of the configuration, DTU events involving only these transcripts were not classified as CDS altering. Contrarily, a lack of known protein domains was still considered as a valid protein domain combination since the absence of domains has functional consequences for the protein. DTU events from or to a configuration where none of the dominant isoforms contained a valid CDS were separately classified as ‘CDS enabling/disabling’ DTU events. All remaining cases were classified as ‘non-CDS altering’. These include for example DTU events without an isoform switch where an isoform without a valid CDS gains or loses dominance or DTU events involving isoforms with CDSs identical to a dominant isoform from the favorite configuration.

CDS data was obtained from AtRTD2 (Zhang et al., 2017). CDSs shorter than 300nt were discarded and isoforms resulting from intron retention events were not considered to contain valid CDSs. Protein domains were determined using InterProScan (v5.22-61, https://github.com/ebi-pf-team/interproscan). All domains from PfamA, TIGRFAM, PIRSF, ProDom, SMART, PrositeProfiles, PrositePatterns, PRINTS, SuperFamily, Coils, and Gene3d were searched, but only domains with a known InterPro identifier were used to determine domain altering DTU. The CDSs and InterPro domains can be found in GTF format in Supplemental Data S4.

The full DTU compendium can be found in Supplemental Data S3. This file contains a matrix of all genes with two or more transcript isoforms (rows) and the 206 public RNA-Seq samples (columns).

### Hierarchical clustering of public RNA-Seq samples based on DTU

The DTU compendium, containing DTU calls for every gene in each of the 206 public samples, was transformed to a binary matrix with only the integers ‘1’ for a DTU event and ‘0’ for no event. Euclidean distances between the columns of this matrix were then used for hierarchical clustering of samples (wards method) with hclust from the stats R package (v3.0.2).

### Gene set enrichment analyses and additional gene features

Gene-GO annotations for all evidence types were downloaded from TAIR (Lamesch et al., 2012) and PLAZA 3.0 (Proost et al., 2015) on 24 January 2017. Data form both databases was concatenated and all parental GO terms were added. Homology data from PLAZA 3.0 Dicots was used to divide genes into five phylostrata (Viridiplantae, Embryophyta, Angiosperms, Eudicots, and Brassicaceae) and to count the number of paralogs for Figure 7. Enrichment analyses were based on hypergeometric tests with a Benjamini-Hochberg correction at FDR=0.05. Only GO and evolutionary gene ages from genes that were expressed in five or more samples and have two or more isoforms were used as background for these tests. Protein-protein interactions (PPI) for Figure 7 were obtained by combining experimental data from CORNET 3.0 (Van Bel and Coppens, 2017) and a recent study using protein arrays (Yazaki et al., 2016).

## ACKNOWLEDGMENTS

This work was supported by the Agency for Innovation by Science and Technology (IWT) in Flanders (predoctoral fellowship to D.V.).

## SUPPORTING INFORMATION

**Supplemental Figure S1. Read number and length of public RNA-Seq datasets.** On October 1st 2015, the Sequence Read Archive was queried for RNASeq experiments for *Arabidopsis thaliana*. 2799 Experiments were found, grouped in 214 studies. 395 Of these experiments contain exclusively paired-end sequenced runs, the other 2404 contain exclusively single-end runs.

**Supplemental Figure S2. Dominance detection optimization.** Two methods were tested for dominant isoform detection: the Percentage of Gene Expression (PGE) method and the FPK Ratio method. For the ratio method, alternatives were tested where the ratio was calculated using the mean, minimum, or maximum instead of the median expression value of the other isoforms. For the PGE method, a version was tested where dominant isoform calls were removed if the difference between the dominant isoform’s PGE and the largest PGE of the non-dominant isoforms was less than 0.05 (delta PGE, performed without iteration). Panel A shows the comparison of the PGE and Ratio method with or without iteration. Panel B shows results of using different thresholds of the Ratio method. Panel C shows parameter optimization for the PGE method. The PGE cutoff was determined with a formula that incorporates the number of isoforms per gene. Red boxes indicate optimal parameters in panels B and C. These benchmarks were performed on simulated datasets of 30 million single-end reads of 100bp. For the PGE cutoff optimization, results are shown for Kallisto alone.

**Supplemental Table S1. Metadata of 206 public RNA-Seq samples**. Short and detailed descriptions for the 206 public RNA-Seq samples. The samples chosen for the RT-PCR validation (Figure 3) and the case studies (Figure 8) are indicated here as well.

**Supplemental Table S2. Simpson et al., 2008 RT-PCR validation.** PS and PSam values for the twenty genes in three organs from Figure 3. For each gene, the identifier suffixes of isoforms with either the distal (dist), proximal (prox), or neither splice events are shown in columns 2-3 respectively.

**Supplemental Table S3. Gene set enrichment analyses for DTU frequencies.** Gene set enrichment statistics are shown for Gene Ontology (GO) terms and phylostrata for non-, rare-, and frequent DTU genes. Hypergeometric tests were used to obtain p-values, which were corrected to q-values with the BenjaminiHochberg method.

**Supplemental Data S1. Kallisto isoform expression values.** This CSV file contains the raw TPM values produced by Kallisto for all isoforms in AtRTD2 (rows) across the 206 public samples (columns).

**Supplemental Data S2. Isoform dominance compendium.** This CSV file contains the isoform dominance calls produced by the ensemble method. 0 And 1 indicate no dominance and dominance respectively for each isoform (rows) in each sample (columns).

**Supplemental Data S3. DTU compendium.** This CSV file contains the DTU calls produced by the ensemble method. The compendium consists of seven possible integer values: -2 indicates no expression for the gene in this sample, -1 indicates that no dominant isoform was detected, 0 indicates that the favorite configuration of dominant isoforms is active, and 1, 2, 3, and 4 indicate non-CDS altering, CDS toggling, CDS altering, and domain altering DTU events respectively. Asterisks indicate isoform switches whereas positive integers without asterisks indicate DTU events without isoform switches.

**Supplemental Data S4. AtRTD2 CDS and protein domains.** This GTF file contains the exons, CDS, and protein domains for each isoform in the AtRTD2 reference transcriptome.

## REFERENCES

Andrews S (2010) FastQC: A quality control tool for high throughput sequence data. In. Babraham Bioinformatics, http://www.bioinformatics.bbsrc.ac.uk/projects/fastqc/

Bray NL, Pimentel H, Melsted P, Pachter L (2016) Near-optimal probabilistic RNA-seq quantification. Nat Biotech 34: 525–527

Brown JW, Simpson CG, Marquez Y, Gadd GM, Barta A, Kalyna M (2015) Lost in Translation: Pitfalls in Deciphering Plant Alternative Splicing Transcripts. Plant Cell 27: 2083–2087

Carrie C, Murcha MW, Millar AH, Smith SM, Whelan J (2007) Nine 3-ketoacylCoA thiolases (KATs) and acetoacetyl-CoA thiolases (ACATs) encoded by five genes in Arabidopsis thaliana are targeted either to peroxisomes or cytosol but not to mitochondria. Plant Mol Biol 63: 97–108

Carvalho RF, Feijao CV, Duque P (2013) On the physiological significance of alternative splicing events in higher plants. Protoplasma 250: 639–650

Chamala S, Feng G, Chavarro C, Barbazuk WB (2015) Genome-wide identification of evolutionarily conserved alternative splicing events in flowering plants. Front Bioeng Biotechnol 3: 33

Cheng CY, Krishnakumar V, Chan A, Thibaud-Nissen F, Schobel S, Town CD (2016) Araport11: a complete reannotation of the Arabidopsis thaliana reference genome. Plant J

Dixon DP, Skipsey M, Grundy NM, Edwards R (2005) Stress-induced protein Sglutathionylation in Arabidopsis. Plant Physiol 138: 2233–2244

Dobin A, Davis CA, Schlesinger F, Drenkow J, Zaleski C, Jha S, Batut P, Chaisson M, Gingeras TR (2013) STAR: ultrafast universal RNA-seq aligner. Bioinformatics 29: 15–21

Engstrom PG, Steijger T, Sipos B, Grant GR, Kahles A, Ratsch G, Goldman N, Hubbard TJ, Harrow J, Guigo R, Bertone P, Consortium R (2013) Systematic evaluation of spliced alignment programs for RNA-seq data. Nat Methods 10: 1185–1191

Erkelenz S, Mueller WF, Evans MS, Busch A, Schoneweis K, Hertel KJ, Schaal H (2013) Position-dependent splicing activation and repression by SR and hnRNP proteins rely on common mechanisms. RNA 19: 96–102

Filichkin S, Priest HD, Megraw M, Mockler TC (2015) Alternative splicing in plants: directing traffic at the crossroads of adaptation and environmental stress. Curr Opin Plant Biol 24: 125–135

Filichkin SA, Priest HD, Givan SA, Shen R, Bryant DW, Fox SE, Wong WK, Mockler TC (2010) Genome-wide mapping of alternative splicing in Arabidopsis thaliana. Genome Res 20: 45–58

Gene Ontology C (2015) Gene Ontology Consortium: going forward. Nucleic Acids Res 43: D1049–1056

Gohring J, Jacak J, Barta A (2014) Imaging of endogenous messenger RNA splice variants in living cells reveals nuclear retention of transcripts inaccessible to nonsense-mediated decay in Arabidopsis. Plant Cell 26: 754–764

Gonzàlez-Porta M, Brazma A (2014) Identification, annotation and visualisation of extreme changes in splicing from RNA-seq experiments with SwitchSeq. bioRxiv

Gonzalez-Porta M, Frankish A, Rung J, Harrow J, Brazma A (2013) Transcriptome analysis of human tissues and cell lines reveals one dominant transcript per gene. Genome Biol 14: R70

Gordon A (2009) FastX Toolkit. In. Hannon Lab, http://hannonlab.cshl.edu/fastx_toolkit/

Griebel T, Zacher B, Ribeca P, Raineri E, Lacroix V, Guigo R, Sammeth M (2012) Modelling and simulating generic RNA-Seq experiments with the flux simulator. Nucleic Acids Res 40: 10073–10083

Guo W, Calixto CP, Brown JW, Zhang R (2017) TSIS: an R package to infer alternative splicing isoform switches form time-series data. Bioinformatics: 1–3

Hayer KE, Pizarro A, Lahens NF, Hogenesch JB, Grant GR (2015) Benchmark analysis of algorithms for determining and quantifying full-length mRNA splice forms from RNA-seq data. Bioinformatics 31: 3938–3945

Hobert O (2008) Gene regulation by transcription factors and microRNAs. Science 319: 1785–1786

Hughes TA (2006) Regulation of gene expression by alternative untranslated regions. Trends Genet 22: 119–122

Iniguez LP, Hernandez G (2017) The Evolutionary Relationship between Alternative Splicing and Gene Duplication. Front Genet 8: 14

Kalyna M, Simpson CG, Syed NH, Lewandowska D, Marquez Y, Kusenda B, Marshall J, Fuller J, Cardle L, McNicol J, Dinh HQ, Barta A, Brown JW (2012) Alternative splicing and nonsense-mediated decay modulate expression of important regulatory genes in Arabidopsis. Nucleic Acids Res 40: 2454–2469

Kanitz A, Gypas F, Gruber AJ, Gruber AR, Martin G, Zavolan M (2015) Comparative assessment of methods for the computational inference of transcript isoform abundance from RNA-seq data. Genome Biol 16: 150

Kelemen O, Convertini P, Zhang Z, Wen Y, Shen M, Falaleeva M, Stamm S (2013) Function of alternative splicing. Gene 514: 1–30

Kim E, Magen A, Ast G (2007) Different levels of alternative splicing among eukaryotes. Nucleic Acids Res 35: 125–131

Klepikova AV, Kasianov AS, Gerasimov ES, Logacheva MD, Penin AA (2016) A high resolution map of the Arabidopsis thaliana developmental transcriptome based on RNA-seq profiling. Plant J 88: 1058–1070

Kodama Y, Shumway M, Leinonen R, International Nucleotide Sequence Database C (2012) The Sequence Read Archive: explosive growth of sequencing data. Nucleic Acids Res 40: D54–56

Lamesch P, Berardini TZ, Li D, Swarbreck D, Wilks C, Sasidharan R, Muller R, Dreher K, Alexander DL, Garcia-Hernandez M, Karthikeyan AS, Lee CH, Nelson WD, Ploetz L, Singh S, Wensel A, Huala E (2012) The Arabidopsis Information Resource (TAIR): improved gene annotation and new tools. Nucleic Acids Res 40: D1202–1210

Li B, Ruotti V, Stewart RM, Thomson JA, Dewey CN (2010) RNA-Seq gene expression estimation with read mapping uncertainty. Bioinformatics 26: 493–500

Li E (2002) Chromatin modification and epigenetic reprogramming in mammalian development. Nat Rev Genet 3: 662–673

Li H, Handsaker B, Wysoker A, Fennell T, Ruan J, Homer N, Marth G, Abecasis G, Durbin R, Genome Project Data Processing S (2009) The Sequence Alignment/Map format and SAMtools. Bioinformatics 25: 2078–2079

Lin H, Ouyang S, Egan A, Nobuta K, Haas BJ, Zhu W, Gu X, Silva JC, Meyers BC, Buell CR (2008) Characterization of paralogous protein families in rice. BMC Plant Biol 8: 18

Marquez Y, Brown JW, Simpson C, Barta A, Kalyna M (2012) Transcriptome survey reveals increased complexity of the alternative splicing landscape in Arabidopsis. Genome Res 22: 1184–1195

Moreau F, Thevenon E, Blanvillain R, Lopez-Vidriero I, Franco-Zorrilla JM, Dumas R, Parcy F, Morel P, Trehin C, Carles CC (2016) The Myb-domain protein ULTRAPETALA1 INTERACTING FACTOR 1 controls floral meristem activities in Arabidopsis. Development 143: 1108–1119

Mortazavi A, Williams BA, McCue K, Schaeffer L, Wold B (2008) Mapping and quantifying mammalian transcriptomes by RNA-Seq. Nat Methods 5: 621–628

Pan Q, Shai O, Lee LJ, Frey BJ, Blencowe BJ (2008) Deep surveying of alternative splicing complexity in the human transcriptome by high-throughput sequencing. Nat Genet 40: 1413–1415

Patro R, Duggal G, Love MI, Irizarry RA, Kingsford C (2017) Salmon provides fast and bias-aware quantification of transcript expression. Nat Methods 14: 417–419

Pertea M, Mount SM, Salzberg SL (2007) A computational survey of candidate exonic splicing enhancer motifs in the model plant Arabidopsis thaliana. BMC Bioinformatics 8: 159

Proost S, Van Bel M, Vaneechoutte D, Van de Peer Y, Inze D, Mueller-Roeber B, Vandepoele K (2015) PLAZA 3.0: an access point for plant comparative genomics. Nucleic Acids Research 43: D974–D981

Reddy AS, Marquez Y, Kalyna M, Barta A (2013) Complexity of the alternative splicing landscape in plants. Plant Cell 25: 3657–3683

Roux J, Robinson-Rechavi M (2011) Age-dependent gain of alternative splice forms and biased duplication explain the relation between splicing and duplication. Genome Res 21: 357–363

Sebestyen E, Zawisza M, Eyras E (2015) Detection of recurrent alternative splicing switches in tumor samples reveals novel signatures of cancer. Nucleic Acids Res 43: 1345–1356

Seo PJ, Kim MJ, Ryu JY, Jeong EY, Park CM (2011) Two splice variants of the IDD14 transcription factor competitively form nonfunctional heterodimers which may regulate starch metabolism. Nat Commun 2: 303

Simpson CG, Fuller J, Maronova M, Kalyna M, Davidson D, McNicol J, Barta A, Brown JW (2008) Monitoring changes in alternative precursor messenger RNA splicing in multiple gene transcripts. Plant J 53: 1035–1048

Staiger D, Brown JW (2013) Alternative splicing at the intersection of biological timing, development, and stress responses. Plant Cell 25: 3640–3656

Sun X, Yang Q, Deng Z, Ye X (2014) Digital inventory of Arabidopsis transcripts revealed by 61 RNA sequencing samples. Plant Physiol 166: 869–878

Thatcher SR, Danilevskaya ON, Meng X, Beatty M, Zastrow-Hayes G, Harris C, Van Allen B, Habben J, Li B (2016) Genome-Wide Analysis of Alternative Splicing during Development and Drought Stress in Maize. Plant Physiol 170: 586–599

Trapnell C, Williams BA, Pertea G, Mortazavi A, Kwan G, van Baren MJ, Salzberg SL, Wold BJ, Pachter L (2010) Transcript assembly and quantification by RNA-Seq reveals unannotated transcripts and isoform switching during cell differentiation. Nat Biotechnol 28: 511–515

Van Bel M, Coppens F (2017) Exploring Plant Co-Expression and Gene-Gene Interactions with CORNET 3.0. Methods Mol Biol 1533: 201–212

Vitulo N, Forcato C, Carpinelli EC, Telatin A, Campagna D, D’Angelo M, Zimbello R, Corso M, Vannozzi A, Bonghi C, Lucchin M, Valle G (2014) A deep survey of alternative splicing in grape reveals changes in the splicing machinery related to tissue, stress condition and genotype. BMC Plant Biol 14: 99

Wang BB, Brendel V (2006) Genomewide comparative analysis of alternative splicing in plants. Proceedings of the National Academy of Sciences of the United States of America 103: 7175–7180

Wiszniewski AA, Bussell JD, Long RL, Smith SM (2014) Knockout of the two evolutionarily conserved peroxisomal 3-ketoacyl-CoA thiolases in Arabidopsis recapitulates the abnormal inflorescence meristem 1 phenotype. J Exp Bot 65: 6723–6733

Wiszniewski AA, Smith SM, Bussell JD (2012) Conservation of two lineages of peroxisomal (Type I) 3-ketoacyl-CoA thiolases in land plants, specialization of the genes in Brassicaceae, and characterization of their expression in Arabidopsis thaliana. J Exp Bot 63: 6093–6103

Yazaki J, Galli M, Kim AY, Nito K, Aleman F, Chang KN, Carvunis AR, Quan R, Nguyen H, Song L, Alvarez JM, Huang SS, Chen H, Ramachandran N, Altmann S, Gutierrez RA, Hill DE, Schroeder JI, Chory J, LaBaer J, Vidal M, Braun P, Ecker JR (2016) Mapping transcription factor interactome networks using HaloTag protein arrays. Proc Natl Acad Sci U S A 113: E4238–4247

Zhang C, Yang H, Yang H (2015) Evolutionary Character of Alternative Splicing in Plants. Bioinform Biol Insights 9: 47–52

Zhang C, Zhang B, Lin LL, Zhao S (2017) Evaluation and comparison of computational tools for RNA-seq isoform quantification. BMC Genomics 18: 583

Zhang R, Calixto CP, Marquez Y, Venhuizen P, Tzioutziou NA, Guo W, Spensley M, Entizne JC, Lewandowska D, Ten Have S, Frei Dit Frey N, Hirt H, James AB, Nimmo HG, Barta A, Kalyna M, Brown JW (2017) A high quality Arabidopsis transcriptome for accurate transcript-level analysis of alternative splicing. Nucleic Acids Res

Zhang R, Calixto CP, Tzioutziou NA, James AB, Simpson CG, Guo W, Marquez Y, Kalyna M, Patro R, Eyras E, Barta A, Nimmo HG, Brown JW (2015) AtRTD - a comprehensive reference transcript dataset resource for accurate quantification of transcript-specific expression in Arabidopsis thaliana. New Phytol 208: 96–101

